# Bayesian inference of power law distributions

**DOI:** 10.1101/664243

**Authors:** Kristina Grigaityte, Gurinder Atwal

**Affiliations:** Watson School of Biological Sciences; Cold Spring Harbor Laboratory, Cold Spring Harbor, NY

**Keywords:** Power Law, Bayesian Inference, Python Package, Jeffreys prior, Mixture Model

## Abstract

Observed data from many research disciplines, ranging from cellular biology to economics, often follow a particular long-tailed distribution known as a power law. Despite the ubiquity of natural power laws, inferring the exact form of the distribution from sampled data remains challenging. The possible presence of multiple generative processes giving rise to an unknown weighted mixture of distinct power law distributions in a single dataset presents additional challenges. We present a probabilistic solution to these issues by developing a Bayesian inference approach, with Markov chain Monte Carlo sampling, to accurately estimate power law exponents, the number of mixtures, and their weights, for both discrete and continuous data. We determine an objective prior distribution that is invariant to reparameterization of parameters, and demonstrate its effectiveness to accurately infer exponents, even in the low sample limit. Finally, we provide a comprehensive and documented software package, written in Python, of our Bayesian inference methodology, freely available at https://github.com/AtwalLab/BayesPowerlaw.

---

Power law distributions abound in empirical data. Examples of where power laws appear include geography (city population sizes (1)), astronomy (moon crater sizes(2)), literature (word usage (3)), biology (T cell clone sizes (4, 5)), networks (connections per node (6)), statistical physics (order parameters in phase transitions (7)), computational neuroscience (power spectra of natural images (8)), and many more. The widespread and enigmatic appearance of these long-tailed distributions has stimulated much research and debate into both their origins and their detection. A number of differing theories have been proposed to explain the generation of power law behavior, providing a useful and simple statistical model of natural observations. Little attention has been paid to mixed models of power law distributions, which may provide a better reduced description of empirical data and suggest multiple generative processes. However, despite our increased theoretical understanding, the accurate detection of actual power laws in both simulated and real world data remains challenging, precisely due to the long-tailed nature of the distributions. Previous inference methods are particularly unsuited in the low sample limit where finite sampling induces greater uncertainty and bias in the estimate, and thus motivates a probabilistic solution rather than a single point estimate. Here, we describe a general and accurate Bayesian formulation of power law inference, providing a posterior distribution of parameter estimates. Furthermore, we derive and implement an objective prior distribution suitable for power law Bayesian inference.

Formally, power law distributions over a variable of interest *x* depend on a single exponent (power) *γ* as shown below,

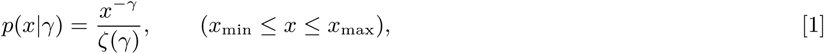

where, in full generality, the power law behavior is assumed to only exist within a certain range of *x* values, (*x*_min_, *x*_max_). The function *ζ*(*γ*) does not depend on *x* and ensures correct normalization of the probability distribution, taking on differing forms depending on whether *x* is a discrete or continuous variable,

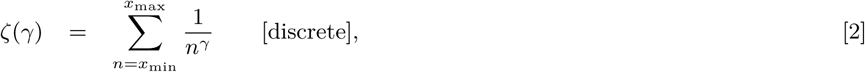

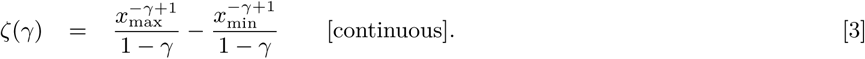

Note that in the discrete case, when *x*_min_ = 1 and *x*_max_ = *∞, ζ*(*γ*) becomes identical to the Riemann zeta function over real values of *γ*. A defining characteristic of power laws is their scale invariance, that is, rescaling the *x* variable by some constant factor *k* leaves the distribution unchanged, up to a normalization factor, *p*(*kx*|*γ*) = *k*^*−γ*^*p*(*x*|*γ*) ∝ *p*(*x*|*γ*). This property underlies the appearance of power law distributions in critical many-body physical systems, leading to the powerful concept of universality and renormalization group in statistical physics and field theory. In critical phenomena the precise value of the exponent indicates membership of a universality class of physical systems, highlighting the need for accurate inference of the exponents.

The simplest and most common method to detect a power law distribution from empirical data is plotting the histogrammed observations on a log-log scale and observing a straight line (Figure 1A,B) (3, 11). Least-square linear regression is then employed on the transformed data to determine the exponent from the slope. Goodness-of-fit tests, such as the Kolmogorov-Smirnov statistic, can be used to indicate how well a power law distribution fits the data (9, 10). Since the data is finitely sampled, the observed straight line can deviate significantly at the tail (3), reflecting the discreteness at low counts. Deviations at either the head or tail of the distribution (Figure 1C,D) can suggest an approximate power law (e.g. *p*(*x*|*γ*) ∝ (1 + *x*)^*−γ*^) or a mixture model of power laws. For example, the human T cell clone size distribution exhibits a heavy tail (Figure 1D), which is likely due to i) sampling noise and ii) substructure stratification of the clones into naive T cells and proliferative non-naive T cells (4).

**Fig. 1.**
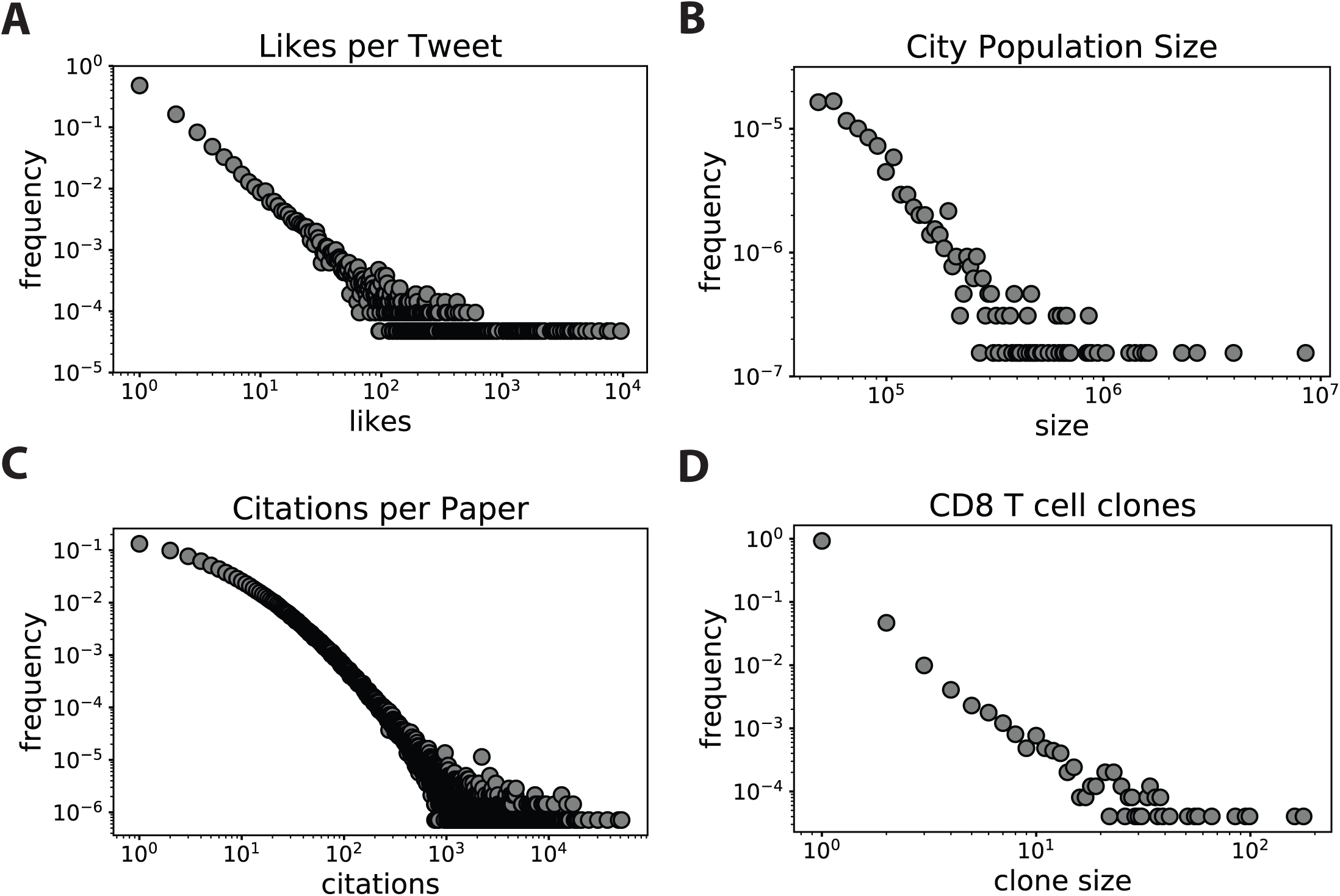
Examples of power law distributions. A) A distribution of likes per tweet - discrete power law. B) A distributions of city population sizes - continuous power law. C) A distribution of citations per paper - an example of the concave power law. D) A distribution of CD8 T cell clone sizes - an example of the convex power law

Despite the widespread use of least-square linear regression on log-transformed data in the literature, this method can result in severe systematic biases in estimates of the exponent (10). This is in part due to the sensitivity of the inference method on the noisy tail end of the empirical distribution, where the data is sparse. Moreover, the traditional assumptions of linear regression no longer hold due to the non-linear log transformation of the assumed Gaussian noise. Thus linear regression, and variations thereof, are both an unreliable and inaccurate estimate of *γ*. An improved method for fitting power laws employs a maximum likelihood (ML) approach (3, 9, 10, 12). While efficient and more accurate ML methods, however, have three important limitations. First, as a point estimate method, ML does not provide a probability distribution over the entire range of probable *γ* values and thus does not equip us with an uncertainty of our estimates. Secondly, ML methods tend to overfit noisy and sparse data, performing poorly in the low sample number regime. Thirdly, they do not allow us to incorporate prior information into our estimates of *γ*. Importantly, we note that all available power law fitting methods implicitly model a single generative process of the data and there are none that can infer a mixed distribution of power laws. In this article we introduce a novel power law fitting approach using Bayesian inference, implemented in a Python software package, that overcomes these limitations imposed by linear regression and maximum likelihood algorithms.

## Bayesian inference with Jeffreys prior

Our goal is to infer the posterior distribution *p*(*γ*|{*x*}) of the exponent variable *γ* given the observed data, {*x*} ≡ *x*_1_, *x*_2_, …, *x*_*n*_,

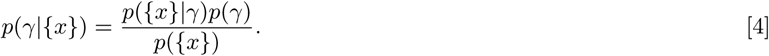

Assuming conditional independence of the data, the likelihood function can be conveniently rewritten as a product, 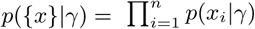. General analytical intractability of the evidence *p*({*x*}) motivates an approximate numerical approach to evaluate the posterior distribution. To this end, we utilize Markov chain Monte Carlo (MCMC) sampling with the Metropolis-Hastings algorithm to closely approximate the desired distribution. Briefly, the MCMC algorithm samples from the posterior distribution by performing a random walk in the parameter space, collecting samples of the exponent *γ* such that the histogram of accepted samples approximates the posterior distribution. We draw potential exponent candidates from a Gaussian distribution and then either accept or reject them by comparing the likelihood of target exponent to the current one. First, we initialize the algorithm by picking an exponent value equal to 1.01 since, natural power law exponents tend to be closer to 1. Next, we perform a burn-in for at least 1000 iterations to diffuse to an an approximately correct value. Normally, the burn-in algorithm is identical to post burn-in sampling with the difference being that values sampled during burn-in do not occur in estimates of the final posterior distribution (13, 14). Here, we modified the burn-in algorithm to speed up the random walk without any observed loss of accuracy. The traditional practice is to reject or accept candidate values of *γ* with some probability, which may lead to accepting a target exponent that is less likely to be correct than the current one (15). During burn-in alone, we do not include this probability and only accept the target exponent if it is more likely to be correct. This allows for a much quicker burn-in, saving time and getting to the right value range in fewer iterations. The burn-in is then followed by the main MCMC in which we do introduce the acceptance probability in the conventional manner.

A longstanding challenge in Bayesian inference is selecting an appropriate prior *p*(*γ*) when *a priori* information is limited. An attractive and principled approach is to use a non-informative prior for the parameter space, also known as Jeffreys prior, that is invariant under monotone transformations (16). Jeffreys prior *p*_*J*_ (*γ*) can be calculated from the Fisher Information, *I*(*γ*),

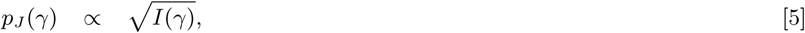

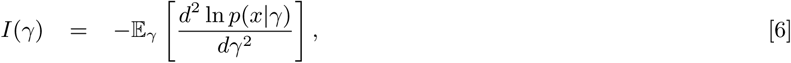

where the expectation in Eq. (6) is taken with respect to the conditional distribution *p*(*x*|*γ*). This prior has been shown to maximize the mutual information between parameters and predictions in the sample number regime where the parameters can be constrained from the data. The prior maximizes the amount of information that can be learned from the data and selects simple effective models in a principled fashion (17). For power law distributions, we find Jeffreys prior to be

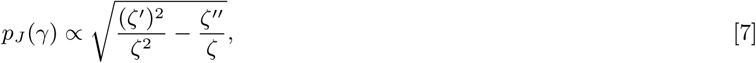

where the derivatives of *ζ* are with respect to *γ*. This formula is correct for both discrete and continuous data, as long as an appropriate *ζ* function is used. In the case of continuous power law distribution where *x*_min_ = 1 and *x*_max_ = *∞* our Jeffreys prior simplifies into equation Eq. (8)

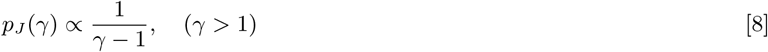

Interestingly, the non-informative prior for a power law Eq. (8) is itself a (truncated) power law, strongly favoring values closer to *γ* = 1, where many natural distributions lie (e.g. Zipf’s law). In our Python software implementation we also enabled the option of a flat prior with variable range, in addition to the default Jeffreys prior.

## Power law mixtures

Historically, when a distribution follows a power law it is assumed that there is only one correct exponent (3, 10, 12). However, heterogeneous datasets, such as complex molecular biological data, are better modeled by a number of generative processes, suggesting a mix of datasets each of which exhibits a power law distribution with a distinct exponent. Fitting such mixture of power law distributions under assumption that only one power law is present will likely lead to inaccurate fit and will fail to recognize the mixture. To infer a mixture model we modify the power law equation Eq. (1) to account for potential mixtures in the data

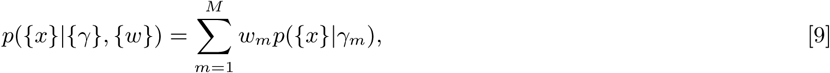

where *M* is number of power laws in the mixture, and {*γ*}, {*w*} are vectors of length *M* containing the exponents and their corresponding weights for each power law in the mixture respectively. This adjustment introduces extra parameters into our previously univariate Bayesian inference equation,

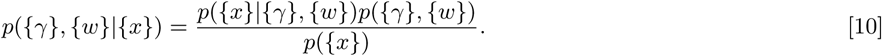

Assuming conditional independence of the data, we obtain the following likelihood function,

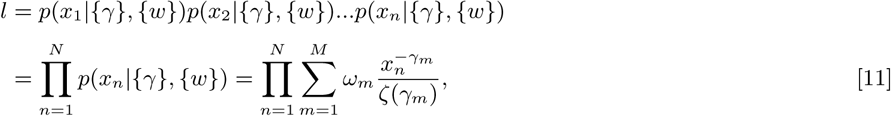

where *N* and *M* are samples size and mixture size respectively. Subsequently, the log likelihood function becomes

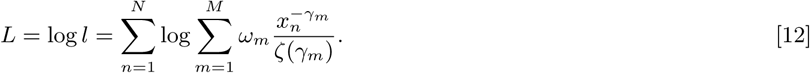

We employed MCMC sampling, as described earlier, to infer the posterior distribution of the set of exponents {*γ*} and the set of weights {*w*}. To determine the optimal number of power laws *M*^*^ that best describes the data we used the Bayesian Information Criterion.

## Results

To demonstrate how our algorithm performs when fitting data sets that follow either a single or two power laws of distinct exponent, we performed simulations where we generated data from both single and mixed power law probability distributions. We show an example of generated data sets of exponents 1.2 (Figure 2A), 2.7 (Figure 2C) and the mixture of the two with weights 0.25 and 0.75 respectively (Figure 2E). We applied our Bayesian inference algorithm to identify the posterior distributions of exponents to see how well our estimated exponent compares to the exponent used for simulations. First, we ran the algorithm assuming that each simulated data set, including the mixed one consisted of only one power law. We found that the algorithm performs well for the data sets that did consist of a single power law (Figure 2B,D), producing a sharply peaked posterior distribution around the true value, but failed to identify any of the exponents from the mixture, instead resulting in a posterior distribution lying between the two correct values (Figure 2F). This implies that fitting a data set under the assumption of only one power law can lead to inaccurate and misleading quantification of the distribution if the distribution consists of more than one power law. Running our Bayesian inference algorithm on the mixed data set of simulated power laws when mixture of two (*M* = 2) is specified resulted in the posterior distribution exhibiting two peaks corresponding to correct exponents (Figure 2G) as well as the weights of each power law distribution in the mixture (Figure 2H).

**Fig. 2.**
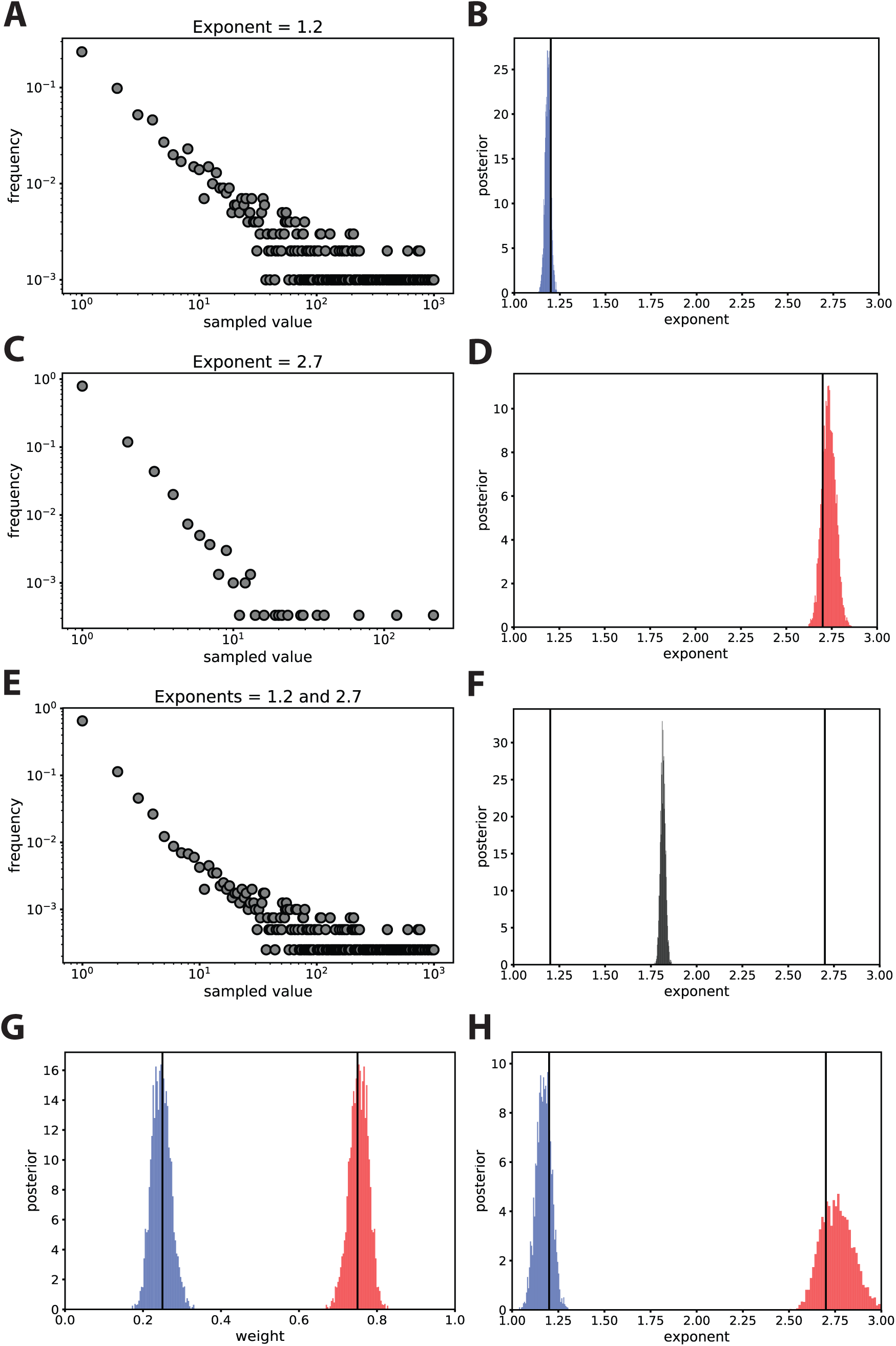
Demonstration of single and mixed power law distribution fits from Bayesian inference with Jeffreys prior. A) Simulated power law distribution of exponent 1.2 and sample size of 1000. B) Posterior distribution of power law exponent in A). C) Simulated power law distribution of exponent 2.7 and sample size of 3000. D) Posterior distribution of power law exponent in C). E) Mixed power law distribution, with 1/4 weight from distribution A) and 3/4 weight from distribution C). F) Posterior distribution of power law exponent in (E) if single power law is assumed. G) Posterior distribution of the two power law weights in the distribution E). H) Posterior distribution of power law (E) exponents assuming the mix of two. Vertical black lines depict correct answers.

### Comparing different fitting methods

To compare our Bayesian inference method to other methods widely used for fitting power law distributions - linear regression-based estimators (11, 18) and maximum likelihood (9, 12) - we performed simulations. Power law distributions of various sample sizes for a range of exponents from one to five were generated and fit using linear regression, maximum likelihood and Bayesian inference (either flat or Jeffreys prior) algorithms (Figure 3). We performed 20 simulations per each exponent and sample size, and calculated the mean and standard deviation across those 20 simulations. In cases of Bayesian inference algorithm, we also compare the mean of posterior distribution means and the mean of the maximum value of the posterior distribution. It is clear that Bayesian inference performs best in all instances with no difference in accuracy between mean and max. Maximum likelihood performs well in the large sample limit, while linear regression trails behind in accuracy amongst all the methods. For all sample sizes - 1000, 100, and even as low as 30 - Bayesian inference performs extremely well. While maximum likelihood is almost as good as Bayesian inference when sample size is large (with an exception of exponents close to 1), it performs poorly when sample size is low such as 30. Linear regression here is done on only the initial part of the distribution (*x*_min_ = 1, *x*_max_ = 10) since otherwise it is completely incorrect due to a heavy and noisy tail. While it performs fairly well for large sample size of exponents in lower range values, it becomes completely inaccurate as sample sizes diminish.

**Fig. 3.**
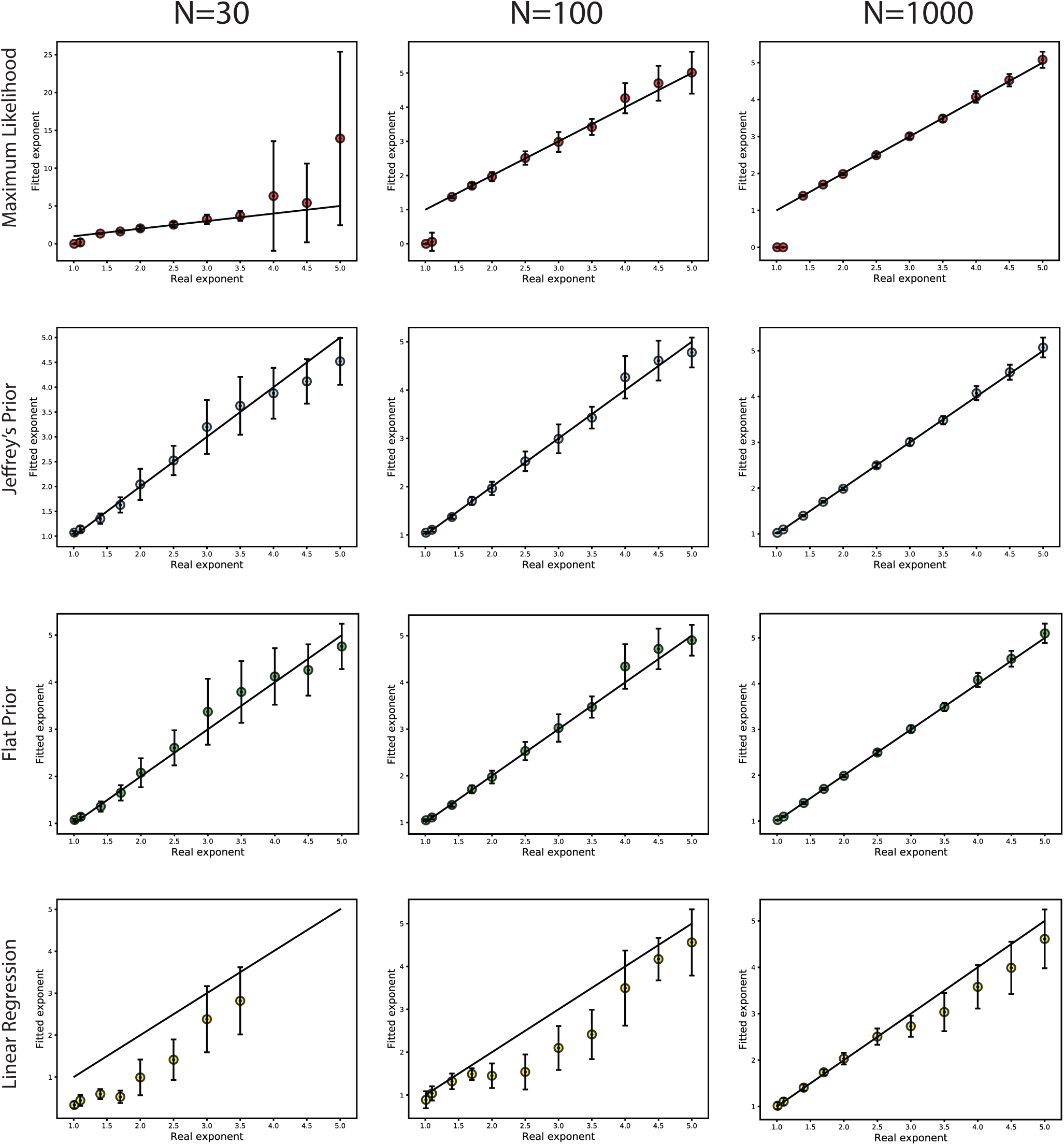
Comparisons of different power law fitting algorithms. 20 power law distributions were simulated per each exponent of samples sizes 30,100 and 1000. Each power law was fitted using either maximum likelihood, Bayesian inference (Jeffreys and flat prior, mean vs max) and linear regression algorithms. Mean estimated exponent and standard deviation of the 20 power laws in each case is plotted against the correct exponent.

### Mixture fitting

To further validate our algorithm with regard to fitting mixtures of two power laws, we performed systematic mixture simulations of various exponent and sample size pairs. We show that the algorithm performs best if (i) the simulated distribution contains a power law with at least one exponent value close to 1 and (ii) if the power law distribution of higher exponent carries larger weight (Figure 4). The reason lower exponent results in better performance of our algorithm is that during the power law simulations there is a higher probability of sampling larger values thus creating a heavier tail. With high exponent, the probability of sampling high values gets lower and the resulting power law is made up of mostly low values. Thus, when combined, the power law with the lower exponent samples a wider range of values, while the power law with the higher exponent only represents a small part of the resulting distribution and is negligible in the tail. Thus, when fitting the mixed distribution, the power law with the lower exponent dominates the algorithm. However, this dominance of the lower exponent gets reduced when the higher exponent power law carries the larger weight. Nevertheless, even if the weighted conditions are not met, our algorithm can still identify the mixture of exponents in most cases, although with the higher uncertainty. The algorithm performs similarly with regard to the weight values as well (Figure 4B).

**Fig. 4.**
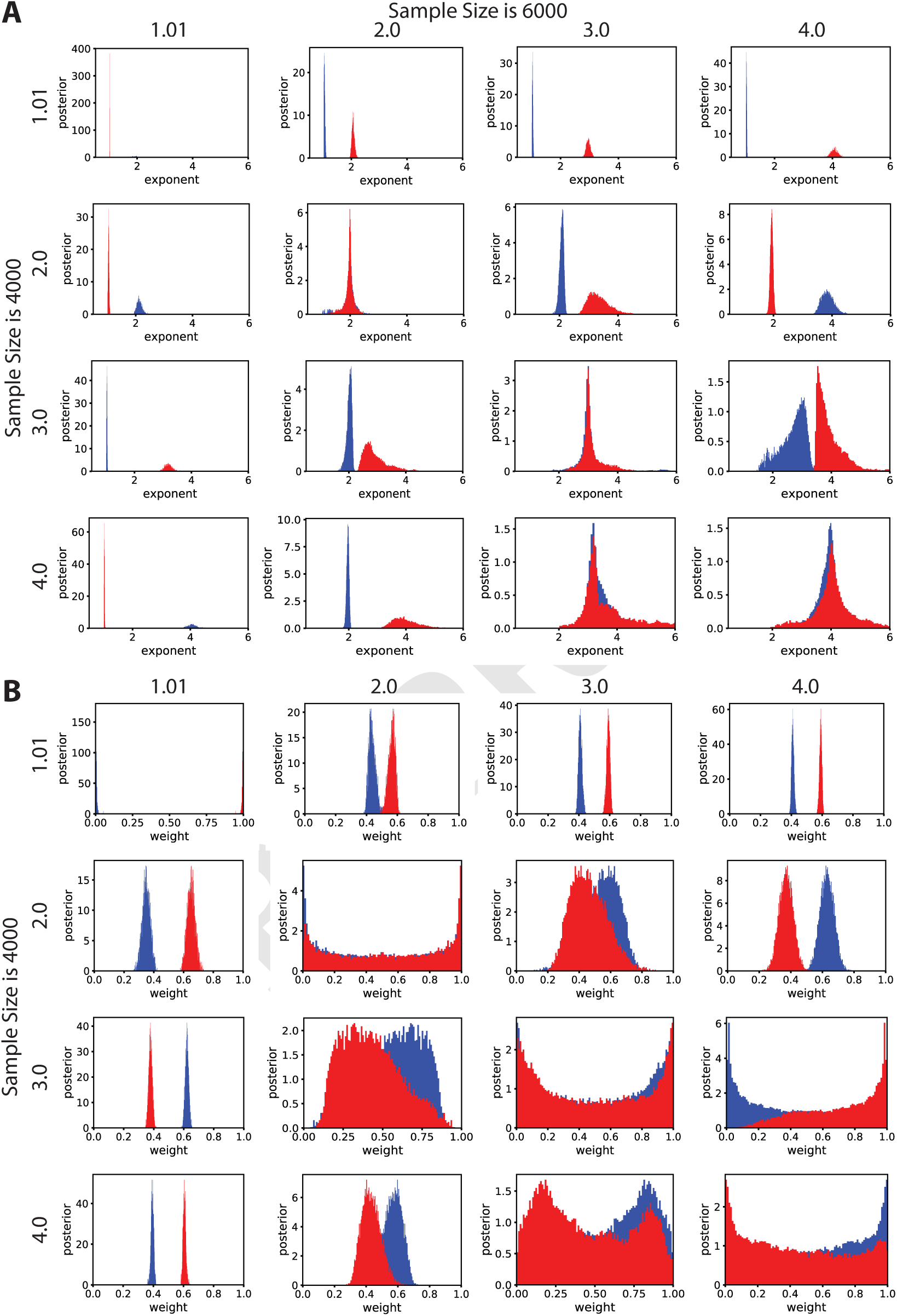
Demonstration of fitting distributions consisting of two power laws. Mixtures of power law distributions were simulated for each exponent pair of samples sizes 4000 and 6000. The generated distributions were fit using our Bayesian inference algorithm to estimate the exponent mixture. A) Posterior distributions of exponents for each power law mixture. B) Posterior distribution of weights for each power law mixture.

### Empirical Datasets

Finally, we used our algorithm to fit power law distributions found in empirical data, some of which have previously identified exponents via different statistical methods (3, 10, 11). Applying our Bayesian inference algorithm on empirical datasets we find that Bayes optimal values of the exponents can sometimes vary significantly from the reported ML results, likely due to the low number of available samples. In this low sample regime the Bayesian method is less sensitive to overfitting due to regularization by the prior distribution. We calculated the Bayes optimal estimates by both the mean of the posterior (mean square error loss) and the mode of the posterior. The Bayes optimal estimate, under mean square error loss, of the empirical power laws varies from 1.03 to 2.44, with relatively small error bars (≈ 2%). The posterior mode closely tracks the posterior mean on real datasets, reflecting the sharply peaked and near symmetric posterior distributions.

We attempted to fit these empirical datasets with mixture models consisting of 1, 2 or 3 power law distributions. The optimal number of mixtures was determined by calculating the Bayesian Information Criteria (BIC). The results on the mixture type, exponents and weights for each data set are shown in Table 1. It is important to note, that the exponents reported here were acquired from fitting the full distribution of each data set, i.e. setting *x*_*min*_ = 1 and *x*_*max*_ = ∞. This ensures that by eliminating some lower bound of data points we did not miss a potential mixture of power laws. We found that only T cell clone size dataset consisted of a mixture of two power laws suggesting two generating mechanisms of distinct T cell subpopulations. Interestingly, this distribution deviates from the straight line on a log-log scale in a convex manner (Figure 1C), consistent with the appearance of the simulated mixture distribution of two power laws with differing exponents (Figure 2E).

**Table 1.** Real dataset fits

| Dataset | Type | Mix | Exponent | Weight | N |
| --- | --- | --- | --- | --- | --- |
| 1. Alice in Wonderland | d | 1 | $1.73 \pm 0.02$ | - | 1465 |
| 2. Lithuanian novel | d | 1 | $1.98 \pm 0.01$ | - | 36896 |
| 3. Harry Potter | d | 1 | $1.63 \pm 0.01$ | - | 5690 |
| 4. Moby Dick | d | 1 | $1.82 \pm 0.01$ | - | 11975 |
| 5. Citations per paper | d | 1 | $1.36 \pm 0.00$ | - | 1401704 |
| 6. Species per genus | d | 1 | $1.67 \pm 0.02$ | - | 1250 |
| 7. Fatalities per terror incident | d | 1 | $1.88 \pm 0.01$ | - | 15531 |
| 8. Injuries per terror incident | d | 1 | $1.51 \pm 0.01$ | - | 12557 |
| 9. Last name usage | c | 1 | $1.81 \pm 0.00$ | - | 151670 |
| 10. Moon crater size | c | 1 | $2.42 \pm 0.02$ | - | 5184 |
| 11. Solar flares | c | 1 | $1.03 \pm 0.00$ | - | 12771 |
| 12. City population size | c | 1 | $2.27 \pm 0.05$ | - | 760 |
| 13. Likes per tweet | d | 1 | $1.73 \pm 0.00$ | - | 20996 |
| 14. All T cell clone size | d | 2 | $2.10 \pm 0.01$ and $4.26 \pm 0.01$ | $0.04 \pm 0.01$ and $0.96 \pm 0.01$ | 173408 |
| 15. CD4 T cell clone size | d | 2 | $1.94 \pm 0.01$ and $4.39 \pm 0.01$ | $0.01 \pm 0.01$ and $0.99 \pm 0.01$ | 66587 |
| 16. CD8 T cell clone size | d | 2 | $2.14 \pm 0.01$ and $4.82 \pm 0.01$ | $0.1 \pm 0.01$ and $0.9 \pm 0.01$ | 24864 |

In addition, we compared the published data set fits acquired using our algorithm to the ones that are published in (3). For a fair comparison, we refit the data with matching *x*_min_ to those reported. The results comparing the exponents are shown in Table 2.

**Table 2.** Real dataset fits

| Dataset | Xmin | Bayes Fit (mean) | Bayes Fit (max) | Maximum Likelihood (Clauset et al, 2009) |
| --- | --- | --- | --- | --- |
| 1. Moby Dick | 7 | 1.96±0.02 | 1.96 | 1.95 |
| 2. Citations per paper | 160 | 2.41±0.01 | 2.40 | 3.16 |
| 3. Species per genus | 4 | 2.07±0.07 | 2.04 | 2.40 |
| 4. Fatalities per terror incident | 12 | 2.39±0.05 | 2.37 | 2.40 |
| 5. Last name usage | 111920 | 2.44±0.10 | 2.41 | 2.50 |
| 6. Solar flares | 323 | 2.22±0.03 | 2.22 | 1.79 |
| 7. City population size | 52460 | 2.34±0.05 | 2.33 | 2.37 |

## Summary

Empirical data from many research disciplines seemingly demonstrate power law behavior, giving rise to a large collection of reported exponents. The varying values of the actual exponents reflects differing mechanistic and generative models underlying the data. Despite the ubiquity of power laws, there has been limited research in accurate inference and measures of uncertainty of the exponents. Here we introduce a novel algorithm for estimating power law distribution exponents using Bayesian inference where the distribution may consist of one or more power law. We derived an objective prior distribution suitable for accurate inference of power law exponents, even in the low sample limit. We demonstrated the accuracy of our algorithm and compared it to previously used linear regression and ML methods with simulated distributions, and show that the Bayesian inference method is superior. Importantly, the Bayesian method provides uncertainty estimates of the inferred parameters, demonstrating increasing confidence as the number of samples increase. On a diverse set of empirical data we find that the Bayes optimal estimates fall within a narrow range close to 1, suggesting a general mechanism and/or giving rise to sampled natural data.

Finally, we provide a comprehensive and documented software package, written in Python, of our Bayesian inference methodology, freely available at https://github.com/AtwalLab/BayesPowerlaw.

## ACKNOWLEDGMENTS

KG was funded by the Ferish-Gerry fellowship from the Watson School of Biological Sciences. GA was funded by the Simons Foundation, Stand Up To Cancer-Breast Cancer Research Foundation Convergence Team Translational Cancer Research Grant, Grant Number SU2C-BCRF 2015-001.

## References

1. Gabaix X (1999) Zipf’s law for cities: An explanation. Q J Econ 114(3):739–767.

2. Cross CA (1966) The size distribution of lunar craters. Mon Not R Astron Soc 134(3):245–252.

3. Newman M (2005) Power laws, pareto distributions and zipf’s law. Contemp Phys 46(5):323–351.

4. Grigaityte K, et al. (2017) Single-cell sequencing reveals *αβ* chain pairing shapes the t cell repertoire. bioRxiv.

5. Bolkhovskaya OV, Zorin DY, Ivanchenko MV (2014) Assessing t cell clonal size distribution: a non-parametric approach. PLoS ONE 9(9):e108658.

6. Barabasi A, Albert R (1999) Emergence of scaling in random networks. Science 286(5439):509–512.

7. Wilson K, Kogut J (1974) The renormalization group and the epsilon-expansion. Physics Reports 12:75–199.

8. Ruderman DL, Bialek W (1994) Statistics of natural images: Scaling in the woods. Physical Review Letters 73:814.

9. Goldstein ML, Morris SA, Yen GG (2004) Problems with fitting to the power-law distribution. Eur. Phys. J. B 41(2):255–258.

10. Clauset A, Shalizi CR, Newman MEJ (2009) Power-law distributions in empirical data. SIAM Rev. 51(4):661–703.

11. Albert R, Barabási AL (2002) Statistical mechanics of complex networks. Rev. Mod. Phys. 74(1):47–97.

12. Alstott J, Bullmore E, Plenz D (2014) Powerlaw: a python package for analysis of heavy-tailed distributions. PLoS ONE 9(1):e85777.

13. Hamra G, MacLehose R, Richardson D (2013) Markov chain monte carlo: an introduction for epidemiologists. Int J Epidemiol 42(2):627–634.

14. van Ravenzwaaij D, Cassey P, Brown SD (2018) A simple introduction to markov chain monte-carlo sampling. Psychon Bull Rev 25(1):143–154.

15. Gilks W (1995) Markov chain monte carlo in practice. (Chapman and Hall/CRC).

16. Gelman A (2009) Bayes, jeffreys, prior distributions and the philosophy of statistics. Stat Sci 24(2):176–178.

17. Mattingly HH, Transtrum MK, Abbott MC, Machta BB (2018) Maximizing the information learned from finite data selects a simple model. PNAS 115(8):1760–1765.

18. Xiao X, White EP, Hooten MB, Durham SL (2011) On the use of log-transformation vs. nonlinear regression for analyzing biological power laws. Ecology 92(10):1887–1894.

